# Sequanix: A Dynamic Graphical Interface for Snakemake Workflows

**DOI:** 10.1101/162701

**Authors:** Dimitri Desvillechabrol, Rachel Legendre, Claire Rioualen, Christiane Bouchier, Jacques van Helden, Sean Kennedy, Thomas Cokelaer

## Abstract

**Summary:** We designed a PyQt graphical user interface – Sequanix – aiming at democratizing the use of Snakemake pipelines. Although the primary goal of Sequanix was to facilitate the execution of NGS Snakemake pipelines available in the Sequana project (http://sequana.readthedocs.io), it can also handle any Snakemake pipelines. Therefore, Sequanix should be useful to all Snakemake developers willing to expose their pipelines to a wider audience.

**Availability:** Source code available on http://github.com/sequana/sequana and standalone on http://bioconda.github.io (sequana package).

## 1 Context and Motivation

Bioinformatics software dealing with biological data (large volumes and variety of data structures) are routinely mingled together to design sophisticated pipeline workflows. Fortuitously, many workflows can be decomposed into *embarrassingly parallel* problems: a task can be applied in parallel on similar data sets (e.g., several DNA samples). Yet, the long-term utility of these pipelines is often tenuous, hampered of interwoven scripts and software libraries, numerous files or complex dependencies between tasks.

Scripting languages may be sufficient to design pipeline workflows. However, they usually lack the ability to handle dependencies between rules, re-entrancies (starting from an intermediate step rather than from scratch), or distributed computing features to handle time-intensive and multi-parametric tasks. In answer to these problems, various workflow managers have emerged as popular tools in the field of bioinformatics (Leipzig,2016). They usually possess all relevant features to design workflows effectively. Consequently, developers have the luxury of choosing a framework amongst many based on personal choice. Influencing factors may include the programming language or the presence of a graphical user interface (GUI). Workflow managers like Galaxy (Goecks,2010) provide GUI drag and drop capability, which can be useful to both developers and end-users. On the other end of the spectrum, command line interface (CLI, hereafter) are still widely employed. Indeed, most developers would prefer a light-weight framework that could be more flexible or accelerate the development of new pipelines.

Amongst the recent CLI-based workflow managers, Snake-make (Köster and Rahmann,2012) has been adopted by a large community of developers. This is especially pronounced in the field of NGS (see http://snakemake.readthedocs.io). Although Snakemake provides a GUI interface, it is server-oriented, which is a limitation on some distributed clusters. Moreover, the GUI is essentially a wrapper of the command line itself; scientists willing to change the behaviour of the pipelines needs to edit the configuration file or masterize the Snakemake arguments. In order to expose Snakemake pipelines to a wider audience, we designed a graphical interface – Sequanix – to offer the ability to edit the configuration file interactively or associate dedicated widgets to parameter types. Sequanix being part of Sequana project (https://sequana.readthedocs.org) was developed to expose Sequana pipelines. However, Sequanix can also handle any Snakemake pipelines as demonstrated in the supplementary data with the SnakeChunks library (Rioualen,2017).

## 2 The graphical interface: Sequanix

*Snakemake* is a text-based workflow that uses Python and a definition language to define rules and workflow properties. A Snakemake pipeline is defined within a file called *Snakefile* (examples in supplementary data). Although not strictly required, pipeline parameters may be externalized within a *configuration* file in YAML or JSON formats.

*Sequana* project provides a set of Snakemake pipelines dedicated to NGS analysis (e.g., RNA-seq, variant calling, …). Hereafter, we distinguish the Snakemake pipelines provided in Sequana from other Snakemake pipelines. We refer to the former as *Sequana pipelines* and to the latter as *Generic pipelines*. The first difference being that every Sequana pipeline is made of a Snakefile *and* a configuration file whereas Generic pipelines may not have a configuration file. The second difference is that the Sequana configuration files are in YAML format only. A third difference is that the configuration file in Sequana must define specific fields which refer to the location or type of inputs files (e.g. FastQ files).

*Sequanix* interface is designed in PyQt, which is a Python binding of the cross-platform GUI toolkit Qt (https://www.qt.io/). The user interface consists of a *main* dialog (see Fig. 1), a *Snakemake* dialog, and a *Preferences* dialog (See supplementary data). The Snakemake dialog is used to configure the Snakemake framework behavior (e.g., number of CPUs to be used) while selection and execution of a pipeline is performed in the main dialog as explained below.

**Fig. 1.**
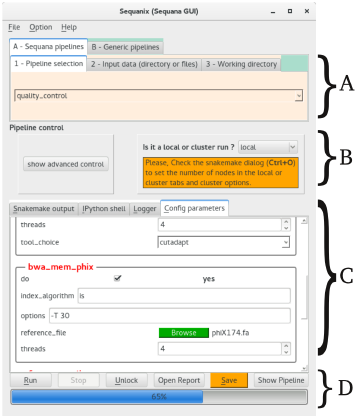
Main Sequanix dialog. One can switch between Sequana and Generic modes or specify the working directory where analysis is performed and results stored (Panel A). The analysis can be run locally or on a distributed computer (Panel B). Configuration file is editable (Panel C). Finally, one can save, run or stop the pipeline execution (Panel D).

At the top of the main dialog (Fig 1, Panel A) one can switch between the *Sequana* or *Generic* mode. In Sequana mode, all released pipelines are shown in *pipeline selection* tab. Once a pipeline is selected, its configuration file is dynamically loaded (Fig. 1, Panel C) and users can modify parameters interactively. Then, users can select the relevant input data (Fig. 1, Panel A). Finally, a working directory must be set where the project (Snakemake pipeline and configuration file edited by the user) is saved. The *Generic* mode works in a similar fashion except that the *input data* tab (specific to Sequana) is now replaced by the *config file* tab. Here, users may provide a configuration file (if needed). The only restriction is that the Snakemake pipeline must be executable in the Sequanix’s environment. In other words, third-party libraries and applications required by the pipeline must be configured properly before starting *Sequanix*.

We specifically designed Sequanix to build upon the Snakemake framework and add important functionalities. For instance, we considered that users should not edit configuration file manually. Therefore, configuration files are loaded in a dedicated widget and their sections and parameters are interpreted and shown in the interface (Fig. 1, Panel C); they can now be edited. Some parameters are associated to specific widgets. For instance a parameter name ending in *_file* or *_browser* becomes a file or directory browser instead of a simple editable line – reducing typing errors. We also propose to annotate configuration file (YAML format) with comments written as Python docstrings (see supplementary). Such comments are then interpreted and appear as tooltips in the GUI.

The Snakemake framework scales without modification, from single and multi-core workstations to cluster engines. This ability reflected in the *Sequanix* interface (Fig. 1, Panel B) and in the Snakemake dialog where one can switch between local and cluster mode, set the number of CPUs, or provide specific job scheduler arguments (e.g., memory requirements).

Once a pipeline (Sequana or Generic) and a working directory are set, the project can be saved (*Save* button) and the pipeline flow (a directed acyclic graph) visualised (*Pipeline* button). Finally, the pipeline can be executed (*Run* Button). Stopping the process (*Stop* button) behaves as a normal Snakemake interruption allowing re-entrancy. If an error occurs (e.g., missing file), one can quickly fix it and re-enter the execution.

## 3 Conclusion

*Sequanix* is a graphical user interface for Snakemake workflows. Users can load, execute and follow the progress of a Snakemake pipeline. The interface is dynamic: new Sequana pipelines are automatically available, configuration files are editable via various widgets, comments in configuration file are interpreted and shown as tooltips. Although Sequanix was primarily developed to expose Sequana pipelines to its end-users, it should benefit a wider community since any Snakemake pipelines can be used. Finally, note that Sequanix is available on Bioconda (https://bioconda.github.io) under the package named **sequana**.

